# Native CRISPR-Cas mediated in situ genome editing reveals the exquisite interplay of resistance mutations in clinical multidrug resistant *Pseudomonas aeruginosa*

**DOI:** 10.1101/496711

**Authors:** Zeling Xu, Ming Li, Yanran Li, Huiluo Cao, Patrick CY Woo, Hua Xiang, Aixin Yan

## Abstract

Antimicrobial resistance (AMR) is imposing a global public health threat. Despite its importance, resistance characterization in the native background of clinically isolated resistant pathogens is frequently hindered by the lack of genome editing tools in these “non-model” strains. *Pseudomonas aeruginosa* is both a prototypical multidrug resistant (MDR) pathogen and a model species for understanding CRISPR-Cas functions. In this study, we report the successful development of the first native type I-F CRISPR-Cas mediated, one-step genome editing technique in a paradigmatic MDR strain PA154197. The technique is readily applicable in additional type I-F CRISPR-containing, clinical/environmental *P. aeruginosa* isolates. A two-step In-Del strategy is further developed to edit genomic locus lacking an effective PAM (protospacer adjacent motif) or within an essential gene, which together principally allows any type of non-lethal genomic manipulations in these strains. Exploiting these powerful techniques, a series of reverse mutations are constructed and the key resistant determinants of the MDR PA154197 are elucidated which include over-production of two multidrug efflux pumps MexAB-OprM and MexEF-OprN, and a typical fluoroquinolone (FQ) resistance mutation T83I in the drug target gene *gyrA*. Characterizing antimicrobial susceptibilities in isogenic strains containing various combinations of single, double, or all three key resistance determinants reveal that i) extensive synergy exists between the target mutation and over-production of efflux pumps, and between the two over-produced tripartite efflux pumps to confer clinically significant FQ resistance; ii) while basal level MexAB-OprM confers resistance only to penicillins, its over-production leads to substantial resistance to all antipseudonmonal β-lactams and additional resistance to FQs; iii) despite the acquisition and over-production of multiple resistant mutations, no obvious evolutionary trade-off of collateral sensitivity is developed in PA154197. Together, these results provide new insights into resistance development in clinical MDR *P. aeruginosa* strains and demonstrate the great potentials of native CRISPR systems in AMR research.

## Author Summary

Genome editing and manipulation can revolutionize the understanding, exploitation, and control of microbial species. Despite the presence of well-established genetic manipulation tools in various model strains, their applicability in the medically, environmentally, and industrially important, “non-model” strains is often hampered owing to the vast diversity of DNA homeostasis in these strains and the cytotoxicity of the heterologous CRISPR-Cas9/Cpf1 systems. Harnessing the native CRISPR-Cas systems broadly distributed in prokaryotes with built-in genome targeting activity presents a promising and effective approach to resolve these obstacles. We explored and exploited this methodology in the prototypical multidrug resistant (MDR) pathogen *P. aeruginosa* by exploiting the most common subtype of the native CRISPR systems in the species. Our successful development of the first type I-F CRISPR-mediated genome editing technique and its subsequent extension to additional clinical and environmental *P. aeruginosa* isolates opens a new avenue to the functional genomics of antimicrobial resistance. As a proof-of-concept, we characterized the resistance development in a paradigmatic MDR *P. aeruginosa* isolate and found extensive resistance synergy and the lack of collateral sensitivity in the strain, implying the extraordinary pathoadaptation capability of clinical MDR strains and challenges to eradicate them.

## Introduction

Antimicrobial resistance (AMR) is imposing an alarming threat to the global public health. Of particular challenging in clinics are those “ESKAPE” pathogens which constitute the major sources of nosocomial infections and are extraordinary to engender resistance, i.e. *<u>E</u>nterococcus spp.*, *<u>S</u>taphylococcus aureus*, *<u>K</u>lebsiella spp.*, *<u>A</u>cinetobacter baumannii*, *<u>P</u>seudomonas aeruginosa*, and *<u>E</u>nterobacter spp*‥ Owing to its intrinsic resistance to a variety of antimicrobials and the enormous capacity of developing acquired resistance during antibiotics chemotherapies, *Pseudomonas aeruginosa* is recognized as the prototypical multidrug resistant (MDR) pathogen [1–4]. Remarkably, in recent years, international high-risk clones of MDR *P. aeruginosa* have emerged and have been shown to cause worldwide outbreaks [5]. Genetic analyses reveal that these clones often contain a complex set of resistance markers [6–8] including both the genetic variations that cause resistance to specific classes of antibiotics, such as mutations in drug targets and acquisition of drug modification enzymes, and those conferring simultaneous resistance to multiple drugs, such as over-expression of multidrug efflux pumps. However, key genetic mutations responsible for the MDR development of clinically significant resistant isolates in their native genetic backgrounds remain undetermined. Furthermore, relative contributions and the interplay of different resistance determinants which shape the MDR profile of resistant isolates and imply the corresponding treatment regimens remain largely elusive.

Current knowledge of the resistance determinants and their mechanisms are largely obtained by reconstitution in laboratory model strains [9, 10]. Several studies indicated that there is a lack of multiplicative or synergetic effects between over-expression of multidrug efflux pumps, and between efflux and the mechanisms causing resistance to specific classes of antibiotics, such as over-expression of the cephalosporinase AmpC and mutations in the DNA gyrase GyrA or topoisomerase IV ParC [11–14]. It was reported that synergistic interactions occur when different types of multidrug efflux systems operate simultaneously, i.e. the tripartite resistance-nodulation-division (RND) efflux pumps and the single component pump (e.g. TetA/C) [15]. Antagonistic interplay of different RND efflux pumps was also reported, especially in the *nfxC* type MDR *P. aeruginosa* isolates in which over-production of the MexEF-OprN pump is often concomitant with an impairment of the MexAB-OprM pump in the strain, resulting in a high resistance to quinolones but a hyper-susceptibility to β-lactams [16, 17]. Compounding the complexity of the resistant mutation networks is the effect of genetic background of resistant strains and epistasis among different resistant mutations which are increasingly recognized to play important roles in resistance development and affect the effectiveness of antibiotic chemotherapies [18–20]. Hence, it is necessary to identify the key resistant determinants and their collateral effects in the native genetic background of clinical MDR strains. Yet, these molecular characterizations are often hindered by the lack of efficient and readily applicable genomic editing tools in these “non-model” strains.

In addition to be a prototypical MDR pathogen, *P. aeruginosa* is an important model system for understating CRISPR-Cas functions, especially the type I CRISPR-Cas system. Phylogenetic analysis revealed that CRISPR-Cas systems are widely distributed in global AMR *P. aeruginosa* isolates with more than 90% belonging to the I-E or I-F subtypes [21]. Owing to its built-in genome targeting activity and reprogrammable feature, in recent years, repurposing the native CRISPR-Cas systems present in the large number of prokaryotes for genetic editing is emerging as a promising strategy for functional genetics, especially in those species with low transformation efficiency and poor homologous recombination. For instance, the native type I-B CRISPR-Cas system in *C. tyrobutyricum*, *C. pasteurianum*, *H. hispanica* (archaea) and the type I-A & III-B native CRISPR-Cas systems in *S. islandicus* (archaea) have been successfully harnessed for genome editing in the corresponding species recently [22–25]. Whether the broadly distributed native CRISPR-Cas systems in *P. aeruginosa*, especially the most common subtype of I-F, can be harnessed for genome editing and functional genomics of antimicrobial resistance remains unexplored.

Previously, we have isolated a MDR *P. aeruginosa* strain PA154197 which displays high epidemic potentials with a resistance profile (resistant to five of the seven commonly used antipseudomonal drugs) comparable to the international high-risk clone ST175 [26]. A large number of resistant mutations were predicted by genomic analysis, including mutations in the DNA gyrase *gyrA* and Cephalosporinase *ampC* which are associated with resistance to fluoroquinolones (FQ) and β-lactams [27, 28], respectively, and gene mutations potentially causing over-production of three multidrug efflux pumps MexAB, MexEF, and MexGHI (S1 Table). Genome sequencing reveals that PA154197 contains a native type I-F CRISPR-Cas locus. These together renders PA154197 a paradigm to explore the harnessing of a native type I-F CRISPR-Cas system for genome editing in a clinical MDR *P. aeruginosa* strain and exploiting the system for AMR characterization in its native genetic background.

In this study, we report the successful development of a single plasmid-mediated, one-step, precise genomic manipulation technique in PA154197 by exploiting its native type I-F CRISPR-Cas system, and a two-step In-Del approach to edit the genomic loci lacking an effective PAM (protospacer adjacent motif) sequence. Exploiting this efficient and powerful technique, a series of single, double, and triple mutations of key resistant determinants are constructed, and their interplay in resistance synergy and potential collateral effects are investigated, providing a comprehensive functional genetics investigation of clinical MDR which have implications in clinical treatment of MDR infections. Lastly, we examine the applicability of the established editing technique in two additional type I-F CRISPR-containing, clinical and environmental *P. aeruginosa* strains PA150567 and Ocean-100. Together, these results demonstrate the general applicability of native CRISPR-based editing system in the characterization of resistance development of clinical MDR *P. aeruginosa* isolates, and presumably other species.

## Results

### PA154197 contains a functional type I-F CRISPR-Cas system

Analysing the genome sequence of PA154197 reveals the presence of the signature *cas8f* and *cas6f* genes and the unique *cas2-cas3* fusion in its CRISPR-Cas loci, suggesting that the system belongs to the subtype I-F (Fig 1A) [29]. The *cas* operon of the system is found to be sandwiched by two convergent CRISPRs. Their consensus repeat sequence differs by only one nucleotide, and their spacers are nearly identical in size (32 bp). In addition, a number of spacers show significant homology to phage or putative prophage sequences (data not shown), suggesting the DNA interference potential of this CRISPR-Cas system and the feasibility of exploiting the system for genome editing.

**Fig 1.**
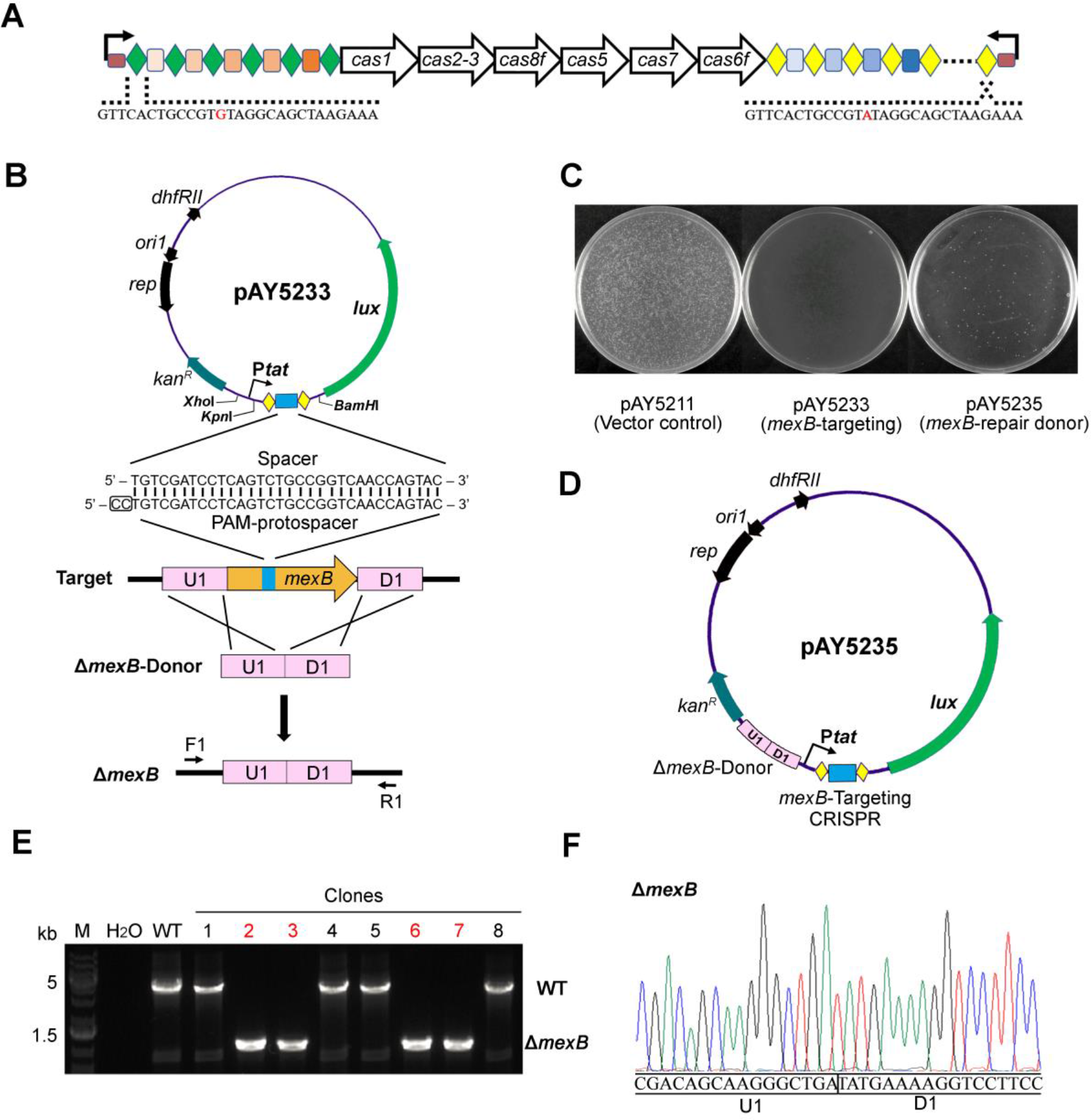
Repurposing the functional native CRISPR-Cas system for gene deletion. **(A)** Schematic representation of the native type I-F CRISPR-Cas in PA154197. Diamonds and rectangles indicate the repeat and spacer units of a CRISPR array, respectively. Curved arrows (in black) above the leader sequence (in brown) indicate the orientation of CRISPR transcription. The consensus repeat of the two CRISPR arrays differs by one nucleotide (in red). **(B)** Schematic showing the design of the *mexB*-targeting plasmid (pAY5233) and the *mexB*-deletion donor. The mini-CRISPR in pAY5233 comprises a 32-bp spacer (in blue) targeting the *mexB* gene and two flanking repeats (in yellow), and is co-expressed with the reporter *lux* operon (in green) under the control of the strong promoter P*tat*. The PAM sequence is framed. The donor (in pink) consists of sequences upstream (U1) and downstream (D1) of *mexB*. The *Kpn*I and *BamH*I sites are used for mini-CRISPR insertion and the *Xho*I site is used for one-step cloning of the donor. **(C)** Representative plates showing the transformation efficiency of the vector control pAY5211, the targeting plasmid pAY5233, and the editing plasmid pAY5235. **(D)** Design of the *mexB*-deletion plasmid pAY5235 which contains both the self-targeting CRISPR and the repair donor. **(E)** Eight randomly selected transformants were subjected to colony PCR to screen for the Δ*mexB* mutants (positive clones are highlighted in red). Primers used in colony PCR (F1/R1) are indicated in panel (B). **(F)** The screened Δ*mexB* mutants in (E) were further validated by DNA sequencing.

To harness the native CRISPR-Cas for genome editing, it is necessary to first examine its endogenous genome targeting activity. We select *mexB* as the target gene for this purpose, which encodes the inner membrane component of the housekeeping efflux system MexAB-OprM. Previous studies revealed that the canonical target of the type I-F CRISPR-Cas system is 5’-CC-protospacer-3’ [30] (note that our study follows the standard guide-centric PAM definition [31]). Hence, an internal 32-bp sequence preceded by a 5’-CC-3’ PAM in *mexB* is selected as the target (PAM-protospacer). A functional mini-CRISPR which is composed of the 32-bp spacer flanked by 28-bp repeat sequences at both ends is then synthesized and cloned into the expression vector pMS402, which contains a kanamycin-resistant gene and a *lux* reporter cassette [32], to generate the targeting plasmid pAY5233 (Fig 1B). To ensure the expression of the mini-CRISPR and efficient targeting, a strong promoter P*tat* [33] is selected and cloned upstream to drive its expression in pAY5233. To overcome the poor antibiotics-based selection of transformants in MDR strains, the P*tat*-mini-CRISPR is positioned upstream of the *lux* operon in frame in pAY5233 such that P*tat* simultaneously drives the expression of both the CRISPR element and the *lux* operon which assists the transformant screening (Fig 1B). When the targeting plasmid pAY5233 and the control plasmid pAY5211 (which contains all the elements described above except the mini-CRISPR fragment) were introduced into PA154197 cells, a dramatic decrease of the transformants recovery was observed comparing to the non-targeting control (Fig 1C), implying the occurrence of the detrimental chromosome cleavage in the pAY5233-containing cells. This result confirms that the native CRISPR-Cas system is active in PA154197.

### Harnessing the native type I-F CRISPR-Cas system to delete the resistance gene *mexB*

To exploit the system for gene deletion (Δ*mexB* as an example), we then assemble a 1-kb donor sequence consisting of the 500-bp upstream and 500-bp downstream of *mexB* for homologous recombination, and insert it into pAY5233 to yield pAY5235, termed as the editing plasmid (Fig 1D). The number of transformants of pAY5235 is significantly increased (by more than 10-fold) compared to pAY5233 (Fig 1C), suggesting the occurrence of homologous recombination by provision of the donor sequence. We randomly selected eight luminescence positive colonies for validation and found four colonies showed the desired, scarless and precise deletion of *mexB* in the chromosome (Fig 1E and 1F). These results demonstrate the success of genome editing by exploitation of the native type I-F CRISPR-Cas system in the clinical MDR isolate PA154197 by one-step introduction of a single editing plasmid.

The success rate of 4/8 seemed moderate. However, we speculate that it was due to the unusually large size (3141 bp) of *mexB* rather than the efficiency of the editing technique, as subsequent attempts to delete shorter fragments, i.e. 50-bp, 500-bp, or 1000-bp within *mexB* (S1A Fig), yielded significantly improved success rate, i.e. 8/8 (50-bp deletion), 7/8 (500-bp deletion), and 7/8 (1000-bp deletion) (S1B Fig). Given that the average length of a prokaryotic gene is ~1 kb [34], the native CRISPR-based editing strategy we developed is efficient to knock out a target gene. Indeed, the 4/8 successful rate for the 3-kb *mexB* is fairly high according to several recent reports [35, 36]. Moreover, we found the editing plasmid can be readily cured following culturing the edited cells in the absence of the antibiotic (kanamycin) pressure overnight (S2 Fig), suggesting the feasibility of multiple rounds of gene editing using the reprogrammable pAY5235 editing platform.

To examine the applicability of the technique in other type I-F CRISPR-Cas containing *P. aeruginosa* strains, we set out to construct *mexB* deletion in a carbapenem resistant clinical strain PA150567 (accession number: LSQQ00000000) isolated from the Queen Mary Hospital, Hong Kong and an environmental strain Ocean-100 (accession number: NMRS00000000) isolated from the North Pacific Ocean [37]. As expected, transformation of the targeting plasmid pAY5233 led to DNA interference in the two strains, and the editing plasmid pAY5235 achieved the desired *mexB* deletion with comparable successful rate as in PA154197 (S3 Fig), confirming the general applicability of the developed editing system in clinically and environmentally isolated, “non-model” *P. aeruginosa* strains.

### Over-expression of the MexAB-OprM and MexEF-OprN efflux pumps contributes substantially to the MDR of PA154197

Three multidrug efflux pumps MexAB-OprM, MexEF-OprN, MexGHI-OpmD were found to be hyper-expressed in PA154197 compared to PAO1 [26]. To test their contribution to the MDR phenotype of the strain, we first delete *mexB*, *mexF*, and *mexH*, which encodes the inner membrane channel of the three efflux systems, respectively. We find that except for IPM, Δ*mexB* leads to substantial decrease in the MICs of all antipseudomonal antibiotics PA154197 is resistant to (Fig 2 and S2 Table), i.e. ATM (64 fold), CAZ (8 fold), TZP (64 fold), MEM (> 32 fold), CAR (>128 fold), LVX (2 fold), and CIP (2 fold), with a greater effect on the antipseudomonal β-lactams (CAR, CAZ, MEM, ATM), and penicillin-β-lactamase inhibitor combinations (TZP) than on fluoroquinolones (LVX and CIP). Δ*mexF* leads to the decrease in MICs of antipseudomonal fluoroquinolones, i.e. LVX (2 fold) and CIP (4 fold), consistent with the previous report of their substrates profiles by ectopic over-expression of the pumps in PAO1 [38]. Double deletion of *mexB* and *mexF* leads to dramatic decrease in MICs of all antipseudomonal antibiotics (except IPM) and several other antimicrobial agents tested (Fig 2 and S2 Table), suggesting that hyper-active drug efflux by these two pumps contributes substantially to the MDR profile of PA154197. Moreover, different from the observations obtained by over-expressing the pumps in *E. coli* which showed no synergy between two tripartite pumps [15], substantial synergy is observed between the over-expressed MexAB-OprM and MexEF-OprN pumps in PA154197 to expel their common substrates FQs as evidenced by the MIC values in the Δ*mexB* Δ*mexF* double deletion strain relative to that in Δ*mexB* and Δ*mexF* single deletion strain. Quantifying these interaction using the fractional inhibitory concentration index (FICI) [39] also demonstrates the substantial synergy of the two pumps in LVX (FICI 0.125) and CIP (FICI 0.188) resistance (Fig 2). Disk diffusion test confirms the susceptibility changes in these strains (S4 Fig). Analysis of the growth curves of these strains in the absence of antibiotics suggests that the observed MIC alterations are not due to intrinsic disadvantages or defects in growth (S5 Fig). Unexpectedly, deletion of another efflux gene *mexH* which is also hyper-expressed in PA154197 (~40-fold higher than in PAO1) [26] does not lead to any detectable difference in the MICs of all the antibiotics and antimicrobial agents tested (Fig 2 and S2 Table). Further deletion of *mexH* in the Δ*mexB* Δ*mexF* strain yields no MIC change either (data not shown), suggesting that it does not contribute to the antibiotic resistance in PA154197.

**Fig 2.**
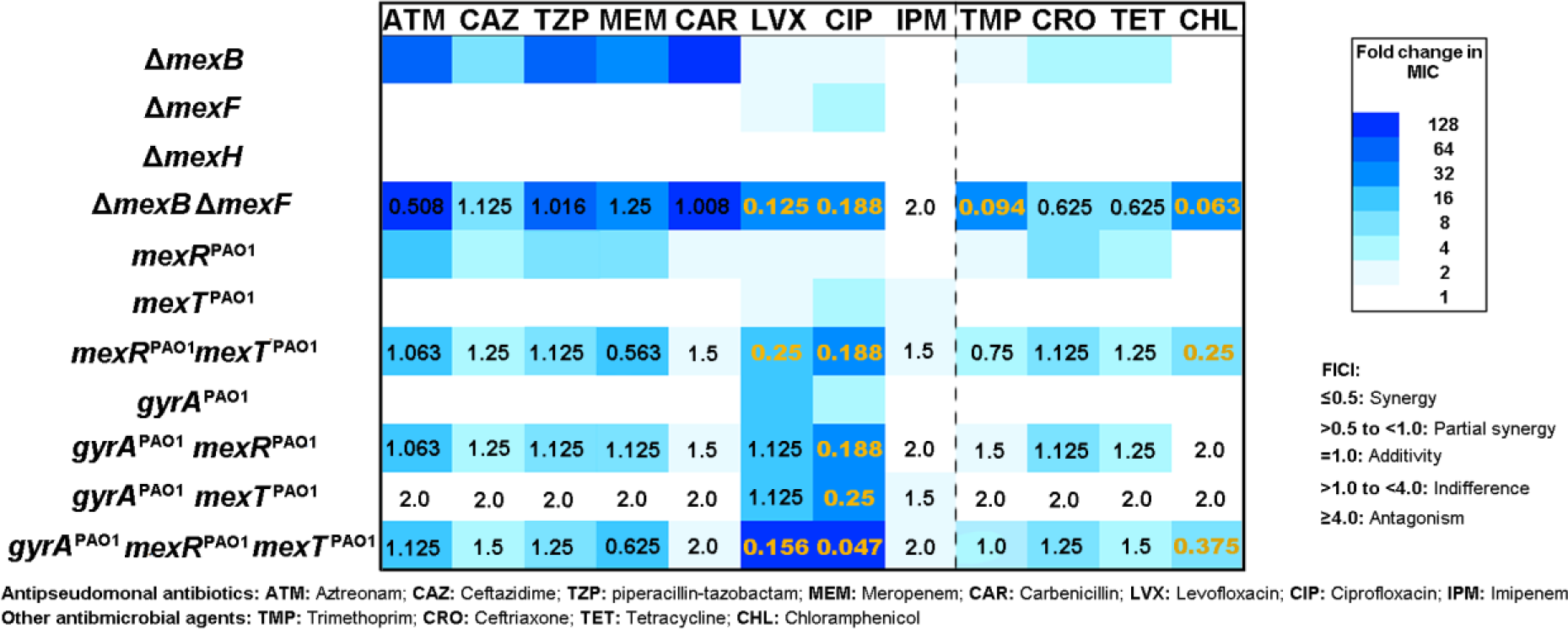
MIC fold-change of various antimicrobial agents in PA154197 isogenic mutants and FICI profiling of reverse mutation combinations. MICs of 12 antimicrobial agents (8 antipseudomonal antibiotics and 4 other antimicrobial agents) were tested in 12 PA154197 isogenic mutants. MIC fold-changes between the wild-type PA154197 and mutants are expressed by the color key shown (detailed MIC values are shown in S2 Table). FICI profiles of various mutation combinations against different antimicrobial agents are presented. FICI values below 0.5 which indicates synergy are highlighted in orange.

### A G226T point mutation in *mexR* is responsible for the *mexAB-oprM* over-expression

Over-expression of efflux genes is often caused by mutations in their transcription regulators. Comparative genomic analysis revealed a single nucleotide substitution G226T (corresponding to introducing a stop codon following E76) in PA154197 *mexR* compared to PAO1 (S6A Fig). Mutations or pre-mature termination of the MexR often leads to over-production of the MexAB-OprM pump (Fig 3A) and MDR in the cells [40–42]. To investigate whether this point mutation accounts for the over-expression of *mexAB-oprM* in PA154197, we reprogramed the editing plasmid pAY5235 by replacing the mini-CRISPR and the donor sequences for deleting *mexB* with those for point mutation of *mexR*, respectively, and constructed the reverse mutation T226G. The resulting construct is designated as *mexR*^PAO1^. Transcriptional level of *mexAB* genes in the *mexR*^PAO1^ mutant is found to be significantly reduced comparing with that in the wild-type (WT) PA154197, to a level that is similar in PAO1 (Fig 3B), suggesting that the G226T mutation in *mexR* is responsible for the over-expression of *mexAB-oprM*. Both MIC analysis and disk diffusion test confirm that *mexR*^PAO1^ causes decreases in MICs of all MexAB-OprM substrates (Fig 2 and S4 Fig).

**Fig 3.**
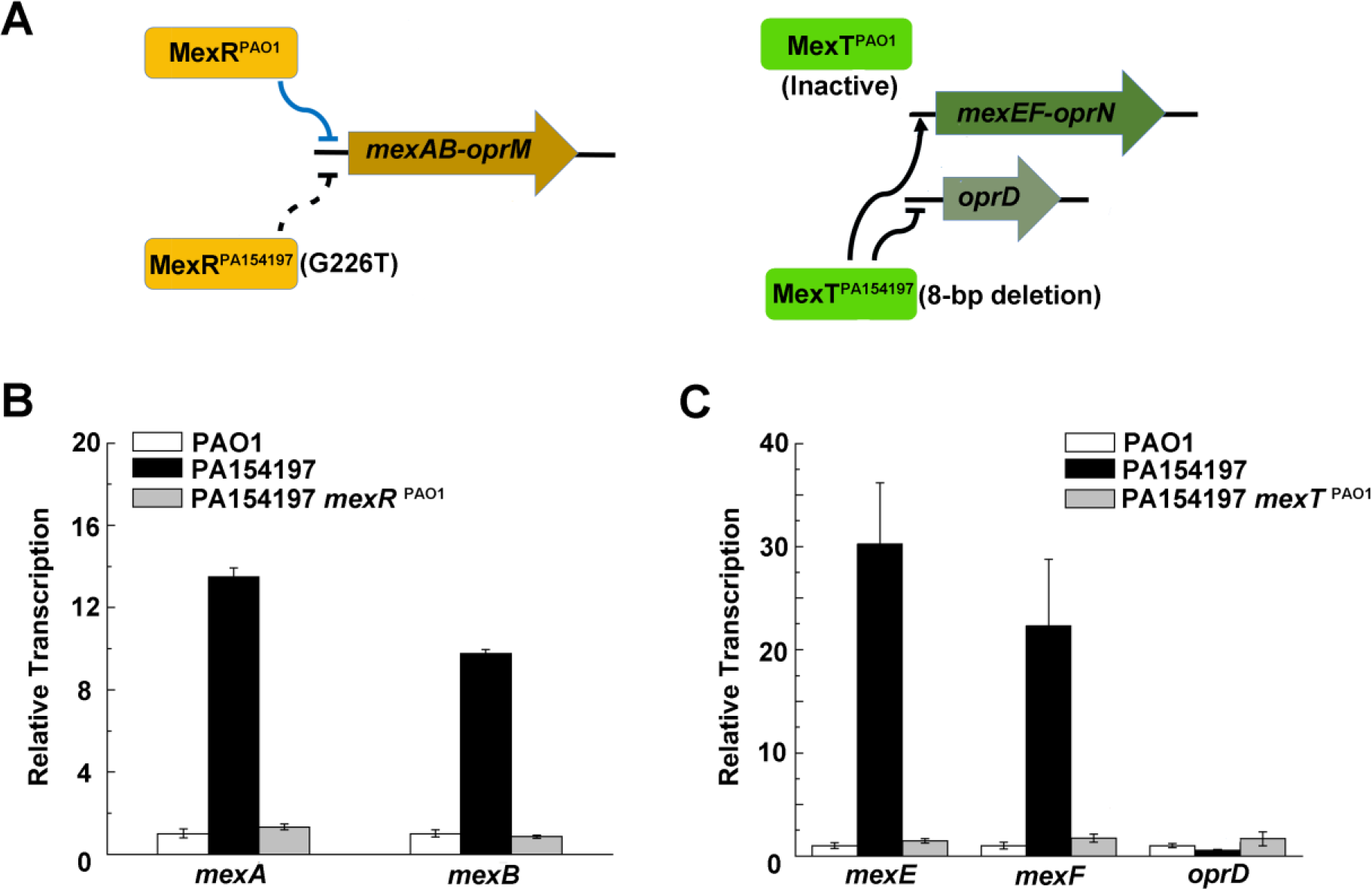
Expression levels of the *mexAB, mexEF* and *oprD* genes in PAO1, PA154197 and its isogenic mutants *mexR^PAO1^ mexT^PAO1^*. **(A)** Schematic showing the regulation of *mexAB-oprM*, *mexEF-oprN* and *oprD* in *P. aeruginosa*. MexR represses the expression of *mexAB-oprM*; MexT activates *mexEF-oprN* and represses *oprD*. **(B)** Transcription alteration of *mexA* and *mexB* in PA154197 and in PAO1, and in PA154197 and its isogenic *mexR*^PAO1^ mutant. **(C)** Transcription alteration of *mexE, mexF* and *oprD* in PA154197 and in PAO1, and in PA154197 and its isogenic *mexT*^PAO1^ mutant.

### An advanced two-step In-Del strategy to edit *mexT* verifies its role in *mexEF-oprN* over-expression and reduced *oprD* expression

The *mexT* gene encodes a transcription regulator of the *mexEF-oprN* efflux system (Fig 3A) [9]. Comparative genomic analysis identified an 8-bp deletion in PA154197 *mexT* (S6B Fig), a *nfxC* type resistant mutation which is proposed to convert the intrinsically inactive, out-of-frame variant of *mexT* as that encoded in PAO1 to a translationally in-framed active MexT that activates *mexEF-oprN* [43, 44]. To verify this effect in the MDR strain PA154197, we attempted to construct the 8-bp insertion reverse mutation. However, construction using the same one-step strategy for gene deletion and point mutation described above was not successful (S7 Fig). Analysing the nucleotides sequence of the PAM-protospacer for the 8-bp insertion reveals the presence of two 6-bp repeats, which potentially obscures the CRISPR recognition of the PAM and the subsequent DNA interference (S7 Fig).

To overcome this limitation, we devise a two-step Insert-Delete (In-Del) strategy to edit this inefficiently targeted site (Fig 4A). The approach bypasses the poorly targeted PAM-protospacer by exploiting a proximal, auxiliary PAM (152-bp downstream of the desired 8-bp insertion site) which can be efficiently targeted to firstly insert a 32-bp short tag (5’- TACAACAAGGACGACGACGACAAGGTGATCAG-3’) between the PAM and the protospacer. The editing of introducing a short tag is applied as both its insertion and subsequent removal is highly efficient. The desired 8-bp insertion is then achieved in the second round of editing during which the short tag is deleted by exploiting the same PAM as in the first step but a different protospacer sequence (termed as the PAM-tag-protospacer) and the provision of a donor sequence which lacks the tag but contains the desired insertion sequence (Fig 4A). The success rate for the first and second round editing for 8-bp insertion in *mexT* is found to be 8/8 and 5/8 (Fig 4B and 4C). This In-Del strategy presumably allows any type of non-lethal genetic manipulations regardless of PAM limitation.

**Fig 4.**
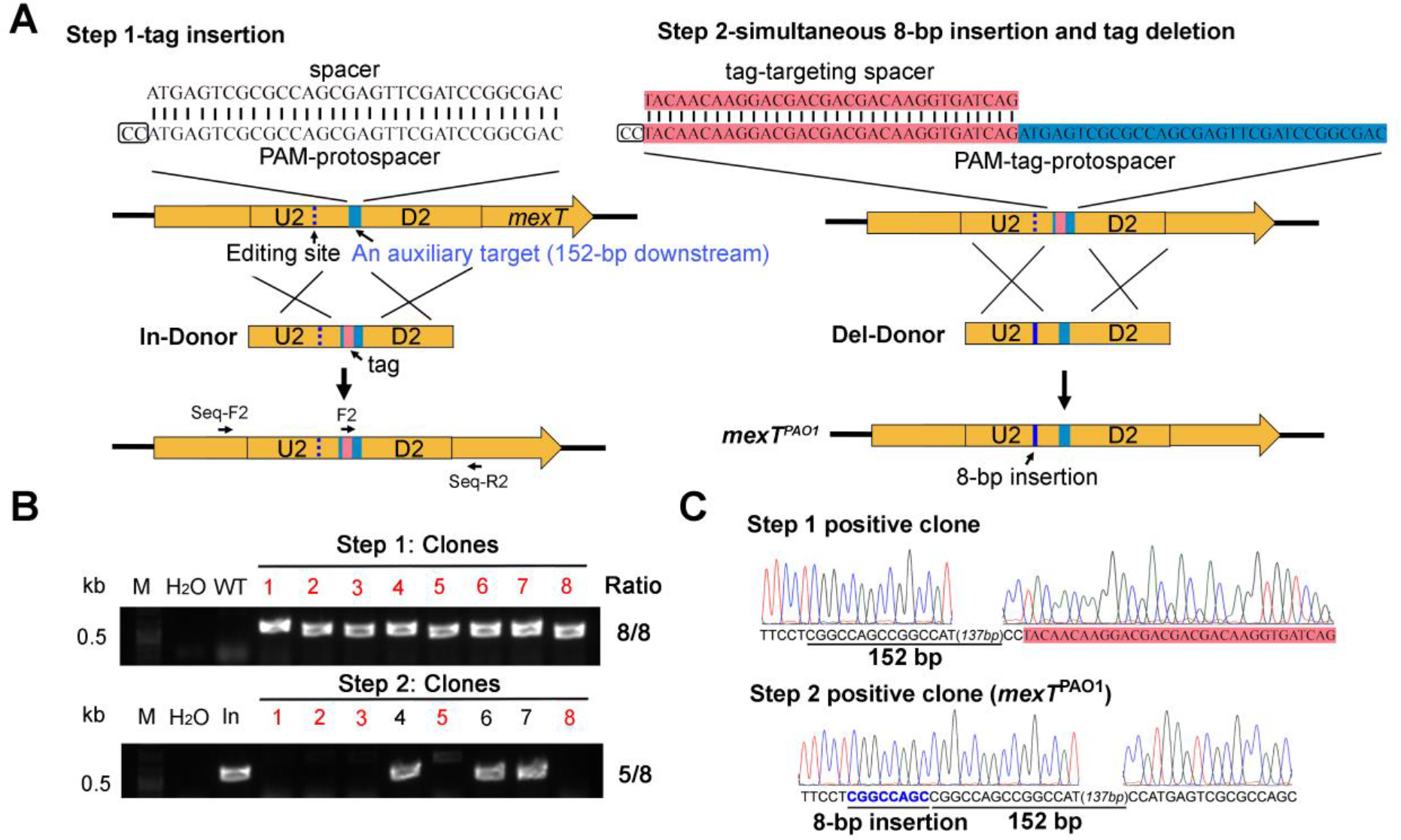
The two-step In-Del editing strategy (taking insertion of 8 bp into *mexT* as an example) **(A)** Schematic design of the two-step In-Del method. In the first step, a 32-bp exogenous DNA sequence (tag, in pink) is introduced to an auxiliary target site (in blue), between its PAM and protospacer portions. In the second step, a tag-targeting CRISPR, as well as a tag-lacking donor which contains the desired mutation (8-bp insertion in this case), is provided to simultaneously remove the exogenous tag and achieve the desired mutation. **(B)** Randomly selected luminescence positive colonies from the two steps were subjected to colony PCR using the primers F2/Seq-R2 (indicated in panel A). Desired mutants are highlighted in red. **(C)** Targets in the potential mutants from the two steps were amplified using the primers Seq-F2/Seq-R2 (indicated in panel A) and validated by DNA sequencing. Representative DNA sequencing results from the two steps are shown, where the tag sequence is shown in pink.

The resulting mutant is designated as *mexT*^PAO1^. As expected, both transcription of *mexEF* and the MICs of LVX and CIP in the *mexT*^PAO1^ cells is reduced comparing with the WT PA154197 (Fig 3C, Fig 2, S2 Table), confirming that the *nfxC* type mutation (8-bp deletion in *mexT*) is responsible for the over-expression of *mexEF-oprN* and contributes to FQ resistance in PA154197. Consistent with the previous report of the *nfxC* type resistant mutation, the reverse mutation *mexT*^PAO1^ also leads to a 3-fold higher transcription of *oprD* than the PA154197 parent (Fig 3C), which encodes a porin protein facilitating the diffusion of carbapenems antibiotics (especially IPM), and an increased susceptibility to IPM in PA154197 (Fig 2 and S4 Fig).

Together, these results indicate that PA154197 may derive from a *nfxC* type resistant strain with acquired resistance to fluoroquinolones and tolerance to IPM (breaking point of IPM is 4). A two-component system ParSR is also known to be involved in regulating *mexEF-oprN* [45]. A nonsynonymous nucleotide substitution G1193A is identified in PA154197 *parS* (S1 Table), However, reverse mutation in this gene does not reveal any susceptibility changes (S2 Table), suggesting this mutation does not contribute to the antibiotic resistance development in PA154197.

### Mutations in the essential gene *gyrA* constitute the third fluoroquinolone resistance determinant in PA154197

Although Δ*mexB* Δ*mexF* leads to a dramatic decrease (32 fold) in MICs of the fluoroquinolones LVX (32 to 1) and CIP (16 to 0.5), the strain still displays a lower susceptibility to these two antibiotics than PAO1 (MIC of both LVX and CIP are 0.25), suggesting the presence of additional FQ resistance determinants in PA154197. A well-studied T248C substitution (corresponding to T83I) in the quinolone resistance determining region (QRDR) which abolishes (fluoro)quinolone binding [46] is identified in the PA154197 DNA gyrase gene *gyrA* (S6C Fig). To verify its role in the high FQ resistance of PA154197 and interaction with the other two FQ resistance determinants identified, we employ the two-step In-Del strategy described for the 8-bp insertion in *mexT* to replace the essential gene *gyrA* in PA154197 with that from PAO1, generating *gyrA*^PAO1^ (S8 Fig). It is found that this gene replacement leads to 8- and 4-fold decrease in the MICs of LVX and CIP, respectively, which is greater than that of Δ*mexB* or Δ*mexF* but not to the susceptible level of PAO1 (Fig 2 and S2 Table), suggesting that the target (*gyrA*) mutation also actively contributes to the FQ resistance in PA154197 and to a greater extent than drug efflux by MexAB-OprM or MexEF-OprN, but it does not mask the contribution of drug efflux.

### Exquisite resistance synergy shapes the MDR profile in PA154197

Previous investigations of resistance mechanisms by reconstitution in the laboratory model strains [10, 38] is often incapable of revealing the relative contribution and synergy of multiple, different resistance determinants owing to its susceptible strain background. The clinical MDR strain PA154197 provides a paradigm to investigate these interactions. We first construct a series of single, double and triple reverse mutations using the efficient editing technique developed and examine the MIC of various antibiotics and antimicrobial agents in these strains (Fig 2 and S2 Table). Our data above have shown that both drug efflux by the over-produced MexAB-OprM and MexEF-OprN pumps and the *gyrA* target mutation contribute to FQ resistance in PA154197 with *gyrA* mutations contributing to a greater extent of resistance than drug efflux. Reverse of all three determinants (*gyrA*^PAO1^ *mexR*^PAO1^ *mexT*^PAO1^ strain) leads to the complete loss of resistance to LVX and CIP, and an overall drug susceptibility in the strain similar to PAO1, indicating these are the key resistant determinants of the MDR PA154197. FICI values (0.156 for LVX and 0.047 for CIP) show that all three determinants synergize the resistance to these two antipseudomonal FQs. Interestingly, MICs in the series of mutants which harbour variable expression levels of efflux systems, i.e. over-production in the PA154197 parent, basal level production in the strains of *mexR*^PAO1^ and *mexT*^PAO1^, and no production in the strains of Δ*mexB* and Δ*mexF*, reveals subtle differences of resistance mechanisms between the two different FQ antibiotics by the interplay of the same set of resistant determinants. For instance, while obvious and extensive synergy is observed in every two combinations of *gyrA* mutation, MexAB over-production, and MexEF over-production for CIP resistance, synergy between target (*gyrA*) mutation and drug efflux to confer LVX resistance is observed when both the MexAB and MexEF pumps are over-produced as evidenced by the FICI values calculated for LVX resistance in *gyrA*^PAO1^*mexR*^PAO1^*mexF*^PAO1^(0.156, synergy), *gyrA*^PAO1^*mexR*^PAO1^ (1.125, indifference), and *gyrA*^PAO1^*mexT* ^PAO1^ (1.125, indifference) (Fig 2). The drug target (*gyrA*) mutation dominates the LVX resistance when only one pump is over-produced.

In addition to FQs, over-production of the MexAB-OprM and MexEF-OprN pumps also synergize the resistance to the non-antipseudomonal antibiotics trimethoprim (TMP) and chloramphenicol (CHL) with an FICI value of 0.094 and 0.063, respectively. Resistance to antipseudomonal β-lactams including the penicillin-β-lactamase inhibitor combination is almost exclusively dependent on the activity of the MexAB-OprM pump in PA154197. Although mediated by the same resistance determinant, the efflux efficiency of different β- lactams by the same MexAB-OprM pump differs, with a greater extrusion efficiency observed for penicillins (CAR), monobactams (ATM), and penicillin-β-lactamase inhibitor combinations (TZP) than for cephalosporins (CAZ) and carbapenems (MEM) (Fig 2 and S3 Table). Notably, MICs of different β-lactams in *mexR*^PAO1^ relative to that in Δ*mexB* cells which reflect the efflux capacity of basal expression of MexAB-OprM (as the level in PAO1) reveals its high capacity of expelling penicillins (CAR) substrate (64 fold) and causing its resistance (S3 Table). MICs in the WT PA154197 relative to isogenic *mexR*^PAO1^ cells which reflects the efflux capacity contributed by over-production of MexAB-OprM reveals its extraordinary capacity of all β-lactams with the greatest extent of resistance enhancement observed in the case of monobactams ATM (16 fold) (S3 Table). The observed FQ resistance by MexAB-OprM is also contributed by its over-production (S3 Table). Combining these, a rather complete substrates profile of the two efflux pumps with relative expel efficiencies in the genetic background of the MDR strain PA154197 is mapped (Fig 5). This information not only advances our understanding of the relative contributions of resistance determinants and their interplay in the clinically isolated MDR strain PA154197 but also has implications in the clinical treatment of drug resistant infections.

**Fig 5.**
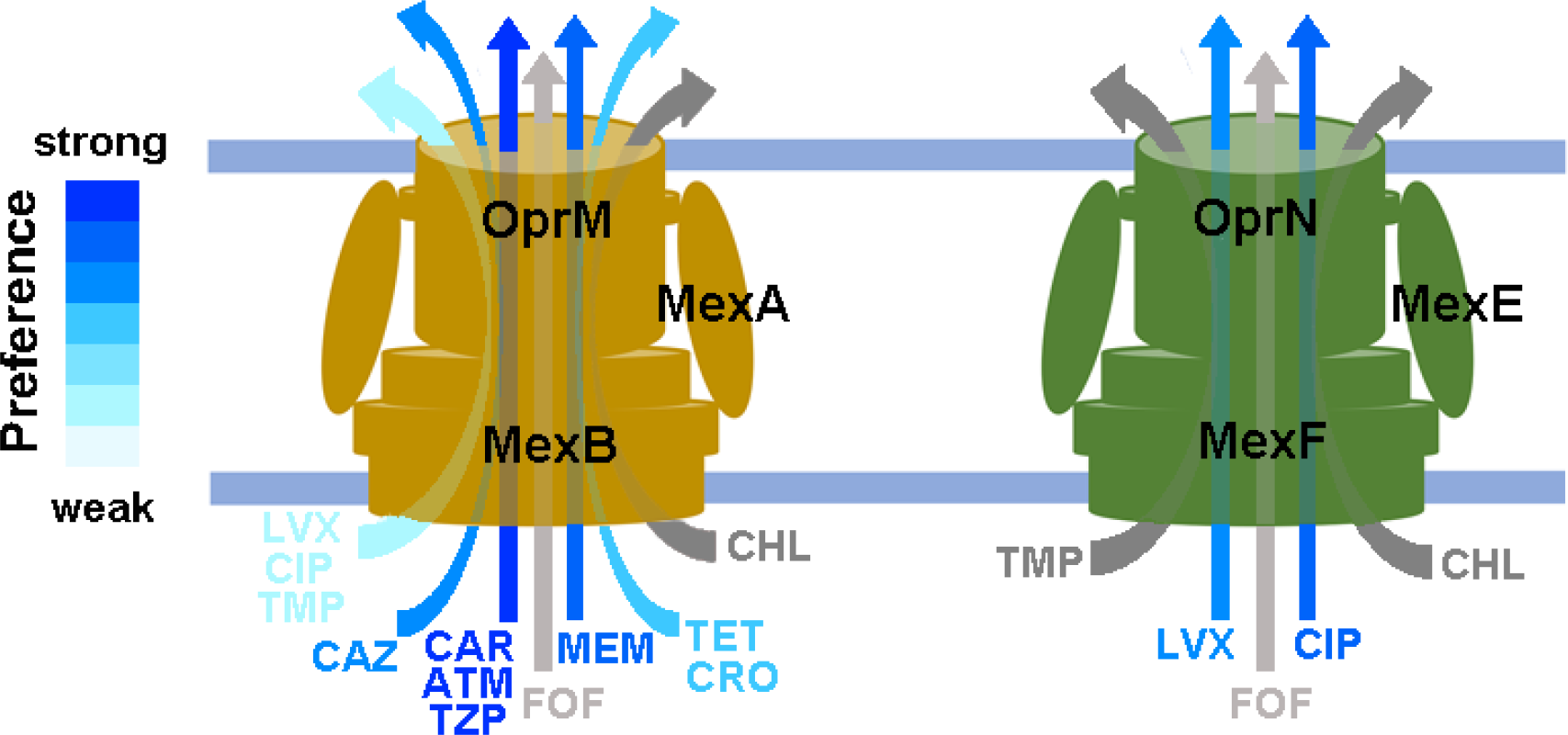
Substrates profile and their relative efflux efficiency by the over-produced MexAB-OprM and MexEF-OprN systems in PA154197. Substrate profiles of the MexAB-OprM and MexEF-OprN and their preferences are shown schematically based on the MIC alterations (Fig 2 and S2 Table) in the Δ*mexB*, Δ*mexF*, and Δ*mexB* Δ*mexF* cells relative to the WT. The greater the MIC changes in the single mutant relative to the WT, the stronger substrates they are denoted. The strength of the substrate preference is expressed by the color key shown, except for the TMP, CHL and FOF (in grey) which MICs were only changed in the double mutant.

### Lacking of collateral sensitivity in PA154197

It was proposed that resistance acquisition often leads to evolutionary trade-offs in resistant strains, i.e. evolved resistance is costly in the absence of the drugs and leads to growth deficiencies relative to the susceptible ancestor; or resistance mutations to one class of antibiotics may exacerbate susceptibility against others, causing a phenomenon called collateral sensitivity. Collateral sensitivity was first described as early as 1950s [47] and was recently verified in a series of laboratory evolved resistant strains of *E. coli* and *P. aeruginosa* [48–50]. It was proposed that collateral sensitivity is a common phenomenon in AMR strains and can be exploited to eradicate clinical resistant strains or reduce resistance development by programming the drug pairs that produce reciprocal collateral sensitivity [39, 51, 52]. However, a recent study reported the lack of collateral sensitivity in clinical *P. aeruginosa* isolates from the cystic fibrosis (CF) patients [53]. But the study did not investigate relative changes in strain susceptibility due to the lack of appropriate baseline controls. The series of strains we constructed which display variable level of resistance in the same genetic background of an MDR strain allows investigating this phenomenon in the context of clinical MDR (a strain from a blood stream infection). To address this, we measure the MICs of the two classes of antibiotics the MDR strain PA154197 displays susceptibility: aminoglycosides (streptomycin and gentamycin) and non-ribosomal peptides (Polymyxin B and colistin) and examine whether the MDR PA154197 is more susceptible to these antibiotics than the series of isogenic, less-resistant “ancestor” strains. However, it is shown that the strains containing various one or two resistant determinants display the same level of susceptibility to aminoglycosides and polymyxins with the WT which contains all three key resistant determinants (S2 Table). Similar result is obtained in the susceptibility to another class of antibiotic which usually is not used to treat *P. aeruginosa* infections, fosfomycin (FOF). These results suggest that unlike those laboratory evolved strains with resistance to one or two specific classes of antibiotics, clinical multidrug resistant strains perhaps have evolved compensatory mutations prevailing the evolutionary trade-off of collateral sensitivity, rendering the treatment of MDR clinical strains especially challenging.

## Discussion

Emergence of resistance to multiple antimicrobial agents in pathogenic bacteria has become a significant global public health threat as there are fewer, or even sometimes no, effective antimicrobial agents to treat the infections caused by these bacteria. The MDR international high-risk clones of *P. aeruginosa* are especially challenging in therapeutics owing to their extraordinary drug resistance and rapid dissemination in hospitals worldwide [54–56]. Previous molecular epidemic analyses have largely focused on identification of genetic variations in resistant isolates in comparison with the model strain PAO1 [57]. To our knowledge, there has been no systematic, functional genetics investigations of resistance development characterized directly in the native genetic background of clinical MDR isolates. In this study, we report the first single-plasmid-mediated, one-step genome editing technique applicable in the native CRISPR-Cas containing, clinical *P. aeruginosa* strains, and its exploitation in MDR characterization which provides new insight into the interplay and collateral effect of resistant mutations in shaping the clinically significant MDR.

In comparison with a recently reported heterologous Cas9-based genome editing method which requires successive transformation of two editing plasmids [35], and the conventional two-step allelic exchange method [58] which takes more than two weeks to construct a mutant with an undesirable FLP site permanently remained in the edited site, our method using a single plasmid to achieve editing in one-step represents a more efficient and clean genome editing technique which can be completed within one week, i.e. 3-4 days for constructing the editing plasmid and 2-3 days for the luminescence-assisted selection and verification. The two-step In-Del strategy further developed to circumvent the limitation of poorly targeted genomic loci principally allows us to conduct any non-lethal genetic manipulation in the bacterial genome (Fig 6), greatly expanding and accelerating the molecular characterizations of the CRISPR-containing, MDR *P. aeruginosa* isolates. The methodology should be readily extended to other clinically significant pathogens, such as *Acinetobacter baumannii* and *Klebsiella spp*. to facilitate resistance characterization and management in these species.

**Fig 6.**
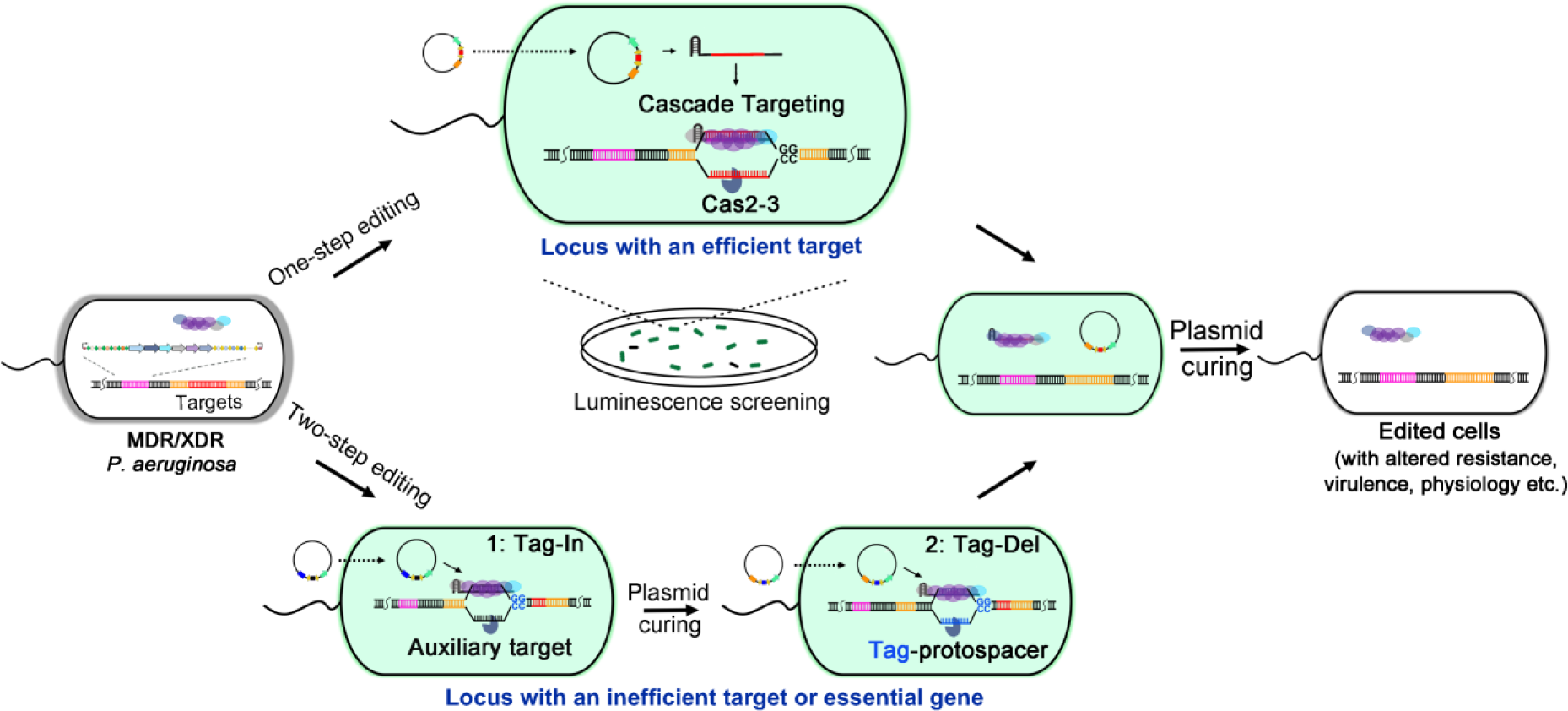
Schematic diagram of the native type I-F CRISPR-Cas mediated genome editing in PA154197 and its exploitations in functional genomics investigation. Desired mutants with altered resistance (or virulence, other physiology) can be obtained in one step by introducing a programmable editing plasmid, which carries a mini-CRISPR (expressing a crRNA) and a repair donor, into the MDR/XDR PA154197 cells with ready luminescence selection. The crRNA directs the Cascade complex to the target. Recombination occurs between the repair donor and the target area to prevent the target interference, resulting in the desired mutation. Two-step editing can be utilized to obtain mutations in the genetic loci that cannot be well targeted by CRISPR. The red sequence indicates a resistant determinant (target) in PA154197. Cells containing the editing plasmid expressing the *lux* genes are shown in green.

Employing this powerful and efficient genome editing methodology, we not only verified the key resistant determinants but also revealed their relative contribution and interplay in shaping the MDR of the clinical strain PA154197 which are unable to be elucidated by reconstitution in the susceptible model strains. The G226T substitution in *mexR* is a newly identified point mutation occurred in PA154197 that leads to the de-repression of *mexAB-oprM*. Three resistant mutations are found to synergize FQ resistance, and mutations in *gyrA* elicit a greater extent of resistance than over-production of MexAB-OprM or MexEF-OprN. We speculate that the G226T mutation in *mexR* (leading to over-production of MexAB-OprM) and the target (*gyrA*) mutations were acquired after the *nfxC* type 8-bp deletion in *mexT* (leading to the over-production of MexEF-OprN and reduced production of OprD) as i) the *nfxC* type resistant strains often contain an impaired *mexAB-oprM* system and additional mutations are presumably needed to achieve simultaneous over-production of both MexEF-OprN and MexAB-OprM pumps; and ii) the contribution of drug efflux (by the MexEF-OprN) to resistance development is not only due to its direct expelling structurally diverse antibiotics but also the capability of driving the acquisition of additional, specific resistance mechanisms by lowering intracellular antibiotic concentration and promoting mutation accumulation [14, 59].

Lastly, we investigated the relative contributions, synergy, and collateral effects of different resistant determinants in conferring the resistance to different classes of antibiotics which should have valuable implications in the treatment of clinical resistant strains. For instance, given the substantial role of drug efflux in causing resistance to CAR, ATM, LVX, and CIP, supplement of pump inhibitors with these antibiotics can conceivably improve the treatment of infections caused by the MDR strain. Notably, studies carried out in a susceptible reference *P. aeruginosa* strain and in *E. coli* showed no synergy between MexAB-OprM and MexEF-OprN in expelling FQs [10]. In contrast to the observation in the laboratory evolved strains [39], no collateral sensitivity and obvious evolution trade-offs such as growth defect in the absence of antibiotics is observed in the clinical MDR strain PA154197. This discrepancy is likely due to the genetic background of the clinical MDR strains and their extraordinary pathoadaptive capabilities remain elucidated. These findings underlie the importance of investigating resistance development in the native genetic background of resistant isolates.

Although genome sequencing and comparative genomics can identify the genetic variations in resistance genes, whether and to what extent these variations contribute to the resistance phenotype cannot be disclosed merely by genomic and transcriptome analyses. For instance, we found that over-expression (~40-fold higher than in PAO1) of the efflux gene *mexH* and nucleotide substitution in *parS* does not contribute to the drug resistance in PA154197. These results suggest that not all genetic variations identified in resistance genes lead to the development of antibiotic resistance, further highlighting the importance of the targeted functional genomics investigations directly in the clinically isolated resistant strains.

## Material and methods

### Bacterial strains, culture conditions

All the bacterial strains used and constructed in this study are listed in S3 Table. *E. coli* DH5α is used for plasmid construction and is usually cultured at 37°C in Luria-Bertani (LB) broth or on the LB agar plate supplemented with 20 μg/ml Kanamycin (KAN). *P. aeruginosa* PA154197 was isolated from the Queen Mary Hospital in Hong Kong, China [26]. PA154197 and its derivatives were selected in LB (broth or agar) with 500 μg/ml KAN at 37°C.

### Plasmid construction

All the plasmids constructed and used in this study are listed in S4 Table. Mini-CRISPR element consisting of two repeats flanking the PAM-protospacer was synthesized by BGI (Shenzhen, China). PCR was performed using the iProof™ High-Fidelity DNA Polymerase (Bio-Rad, USA). Mini-CRISPR elements and plasmid pAY5211 were digested using the restriction enzymes KpnI and BamHI (NEB, USA) and ligated using the Quick Ligation Kit (NEB, USA) to generate the targeting plasmid. Donor sequences which typically contain 500-bp upstream and 500-bp downstream of the editing sites were amplified by PCR and ligated into the linearized targeting plasmid (digested by XhoI (NEB, USA)) using the ClonExpress One Step Cloning Kit (Vazyme, China). All the constructed plasmids were verified by DNA sequencing (BGI, China).

### Transformation of PA154197

Electrocompetent PA154197 cells were prepared by firstly inoculating a fresh colony in LB broth and grown at 37°C overnight with 220-rpm agitation. Following subculture into 50 ml fresh LB broth and growing to OD_600_ = 0.5, cells were collected by centrifugation and washed three times with cold autoclaved Milli-Q H_2_O. The resulting cells were resuspended into 1 ml Milli-Q H_2_O. 100 μl electrocompetent PA154197 cells were then mixed with 1 μg editing plasmid and subject to electroporation (BTX, USA). 1 ml cold LB broth was added to recover the cells. Following culturing at 37°C for 1 h with agitation, cells were pelleted and resuspended in 100 μl LB for spreading (LB+KAN500). Transformants colonies were obtained after incubation at 37°C for 16-20 h.

### Mutant screening and verification

Colonies were firstly subjected to luminescence screening using the Synergy HTX Plate Reader (Bio Tek, USA). Colonies with high luminescent intensity were further verified by colony PCR using Taq DNA polymerase (Thermal Scientific, USA) with indicated primers and DNA sequencing (BGI, China). Sequencing results were visualized using DNA sequencing software Chromas (Technelysium Pty Ltd, Australia)

### Curing of editing plasmid

*P. aeruginosa* cells underwent one round of editing was streaked onto the LB ager plate and incubated at 37°C overnight. Single colony was selected and the curing was verified by the failure of growth in LB with 100 μg/ml KAN.

### Minimum inhibitory concentration (MIC) measurement

MIC was measured following the standard protocol of ASM with slight modification [60]. A single fresh colony of PA154197 was inoculated in LB medium overnight at 37°C with agitation. Overnight culture was then diluted to cell density of 10^5^/ml, and 200 μl of the cells were distributed to each well of the 96-well plate. Antibiotics were then added with final concentrations ranging from 0.25 to 128 μg/ ml. Plates were incubated at 37°C for 16-20 h and MIC value was determined by absorbance at 600 nm.

### Reverse transcription (RT)-quantitative PCR (qPCR)

Bacterial cells from overnight culture were harvested by centrifugation at 4°C. Total RNA was extracted using RNeasy Mini Kit (Qiagen, Germany) according to the manufacturer’s instruction. Reverse transcription was performed using PrimeScript RT reagent Kit (Takara, Japan). qPCR was performed using specific primers and the SYBR Green PCR master mix (Applied Biosystems, USA) in a 20 μl reaction system. The reaction was performed in ABI StepOnePlus real time PCR system with *recA* and *clpX* as reference genes to normalize the relative expression of the target genes. The results were expressed as fold change of the expression of target genes, and results were presented as the mean of three independent biological isolates.

### Disk diffusion assay

20 μl overnight culture as described above was mixed with 5 ml melted LB top agar (0.75 %) and poured on a LB agar plate. After the agar was solidified, round filter paper disks were placed. 5μl antibiotic solution (2 mg/ml) was added to the centre of the paper disks. The plates were incubated at 37°C for 16 h.

### Fractional inhibitory concentration index (FICI) determination

Synergy of mutation combinations was determined using the FICI method [39]. The MIC of the mutant with combined mutations is divided by the MICs of the mutants with single mutation, yielding the fractional contribution of each mutation in the combination. Interaction of different mutations (A, B, C) is scored using the following formula: FICI_(ABC)_=MIC_(ABC)_/MIC_(A)_+ MIC_(ABC)_/MIC_(B)_+ MIC_(ABC)_/MIC_(C)_.

## Acknowledgements

We thank Prof. Susumu Yoshizawa and Prof. Kazuhiro Kogure (Both from the University of Tokyo) for sharing the ocean-100 strain. We appreciate Dr. Xin Deng (Department of Biomedical Sciences, City University of Hong Kong) for the pMS402 vector and Dr. Zhaoxun Liang (School of Biological Sciences, Nanyan Technological University, Singapore) for his advice on the manuscript writing.

## Supporting information

**S1 Fig. Effect of desired truncation lengths on the efficiency of editing. (A)** Schematic diagram of the genetic locus of *mexB* and adjacent region. Desired truncation fragments of 50 bp (B1, 74-123), 500 bp (B2, 74-573), 1000 bp (B3, 74-1073), the full length (*mexB*, 1-3141), and the primers to test these truncations are shown. **(B)** Positive clones of each of truncations in eight randomly selected transformants are highlighted in red.

**S2 Fig. Curing of the editing plasmid in the edited cells.** Edited cells could not grow in the presence of kanamycin (100 μg/ml) after plasmid curing, which was achieved by culturing the edited cells without antibiotic pressure overnight.

**S3 Fig. Harnessing the native type I-F CRISPR-Cas system for genome editing in other *P. aeruginosa* isolates.** The single editing plasmid pAY5235 was transformed in the clinical *P. aeruginosa* isolate PA150567 and the environmental isolate Ocean-100 to achieve *mexB* deletion in one-step. Eight randomly selected luminescent transformants were subjected to colony PCR to screen for the desired Δ*mexB* mutants. Positive clones are highlighted in red. Primers used in colony PCR are indicated in Figure 1B.

**S4 Fig. Disk diffusion assay of the antibiotic susceptibilities of PA154197 and its isogenic mutants.** Disk diffusion test is conducted to examine the susceptibility of PA154197 and its isogenic mutants to eight antipseudomonal drugs **(A)** and four other antibiotics and antimicrobial agents **(B)**. Larger inhibition zone represents the higher susceptibility to the indicated antibiotics.

**S5 Fig. Growth curves of PA154197 and its isogenic mutants in the absence of antibiotics.** No difference of growth among PA154197 and its isogenic mutants were observed.

**S6 Fig. Depiction of genetic mutations identified in the key antibiotic resistance genes *mexR*, *mexT* and GyrA in PA154197. (A)** The point mutation (G226T) in PA154197 *mexR* gene leads to premature termination of its protein product. **(B)** An 8-bp deletion in PA154197 *mexT* relative to that in PAO1 converts the out-of-frame *mexT* product to the in-frame, active form of *mexT* product. **(C)** PA154197 *gyrA* gene contains a T248C substitution (corresponding to T83I in the GyrA protein) in the quinolone resistance-determining region (QRDR) and a ^2723^CCGAGT^2728^ 6-bp deletion (corresponding to the deletion of E959&S960) in the C terminal domain of the gene product.

**S7 Fig. Non-efficient targeting at the desired editing site in the *mexT* gene. (A)** Schematic showing the desired 8-bp insertion (CGGCCAGC) into the *mexT* gene. The 6-bp repeat sequences (in red) in the target which potentially obscure the targeting by the CRISPR-Cas apparatus are shown. PAM is shown in frame. **(B)** No desirable DNA interference was observed by transforming the *mexT*-targeting plasmid pAY5900 into PA154197 cells in comparison with the control plasmid pAY5211. Note that no repair donor was provided in this plasmid.

**S8 Fig. Replacing the essential gene *gyrA* with that from PAO1 using the two-step In-Del method. (A)** Schematic showing the two-step strategy for *gyrA* gene replacement. The tag sequence (in pink) is first introduced downstream of *gyrA* (in orange) between the PAM and protospacer portions of a selected target (in blue). In the second step, a tag-targeting plasmid carrying the repair donor (Del-donor) is provided. Del-donor lacks the tag sequence but contains the *gyrA* gene (in dark blue) from PAO1, which is flanked by two ~1000-bp sequences upstream and downstream of the PA154197 *gyrA* gene. The primer pairs F/Seq-R and Seq-F/Seq-R were used for the colony PCR analysis in (B) and DNA sequencing, respectively. **(B)** Eight luminescent positive transformants from each editing step were randomly selected for colony PCR analysis. Desired mutants confirmed by DNA sequencing are highlighted in red.

**S1 Table. A summary of mutational changes in the antibiotic resistance (AR) genes in PA154197 by comparative genomic analysis**

**S2 Table. MICs of various antibiotics in PAO1, PA154197 and its isogenic mutants**

**S3 Table. Contribution of basal-level, over-produced and overall MexAB-OprM pump to the resistance to different antipseudomonal antibiotics in PA154197**

**S4 Table. Bacterial strains, plasmids used in this study**

