## Supplementary material for "Native CRISPR-Cas mediated in situ genome editing reveals the exquisite interplay of resistance mutations in clinical multidrug resistant *Pseudomonas aeruginosa*": S1 Table

| **Table S1.** A summary of mutational changes in the antibiotic resistance (AR) genes in PA154197 by comparative genomic analysis | | | |
| --- | --- | --- | --- |
| **AR genes** | **Contribution to AR** | **Mutation(s)** | **Reference** |
| *ampC* | Cephalosporinases, cephalosporins resistance | T105A | [1] |
| *gyrA* | DNA gyrase subunit A, (fluoro)quinolone resistance | T248C (T83I in GyrA), 6-bp deletion | [2] |
| *mexR* | Repressor of *mexAB-oprM* , MDR to the substrates of MexAB-OprM | G226T (E76*) | - |
| *armR* | Anti-repressor of *mexR*, MDR to the substrates of MexAB-OprM | G62C, C84T, A95G, A117G | - |
| *nalC* | Repressor of *armR*, MDR to the substrates of MexAB-OprM | G212A, G458C, A625C | [3] |
| *parS* | Two-component regulatory system of *mexEF-oprN*, MDR to the substrates of MexEF-OprN | G1193A | [4] |
| *mexT* | Activator of *mexEF-oprN* efflux system, MDR to the substrates of MexEF-OprN | 8-bp deletion | [5] |
| *oprD* | Porin, entry of IPM | 21 point mutations, 6-bp deletion | - |
| *mexGHI-*  *opmD* | MexGHI-OpmD efflux system | G7C, G80C, C134T, T146C, C218A,  in P*mexGHI-opmD* | - |
| New mutations identified in PA154197 are highlighted in red. *: Stop codon | | | |
