## Supplementary material for "Native CRISPR-Cas mediated in situ genome editing reveals the exquisite interplay of resistance mutations in clinical multidrug resistant *Pseudomonas aeruginosa*": S2 Table

| **Table S2:** MICs of various antibiotics in PAO1, PA154197 and its isogenic mutants | | | | | | | | | | | | | | | | | |
| --- | --- | --- | --- | --- | --- | --- | --- | --- | --- | --- | --- | --- | --- | --- | --- | --- | --- |
| **Strains** | **ATM** | **CAZ** | **TZP** | **MEM** | **CAR** | **LVX** | **CIP** | **IPM** | **TMP** | **CRO** | **TET** | **CHL** | **STR** | **GEN** | **FOF** | **PMB** | **CST** |
| PAO1 | 4 | 1 | 2 | 0.5 | 32 | 0.25 | 0.25 | 2 | 32 | 4 | 8 | 32 | 8 | 2 | 32 | 1 | 0.5 |
| PA154197 | **64** | **16** | **32** | **4** | **>128** | **32** | **16** | **4** | **128** | **64** | **32** | **>128** | **16** | 2 | 32 | 1 | 0.5 |
| Δ*mexB* | 1 | 2 | 0.5 | <0.125 | 1 | 16 | 8 | 4 | 64 | 16 | 8 | >128 | 16 | 2 | 32 | 1 | 0.5 |
| Δ*mexF* | 64 | 16 | 32 | 4 | >128 | 16 | 4 | 4 | 128 | 64 | 32 | >128 | 16 | 2 | 32 | 1 | 0.5 |
| Δ*mexH* | 64 | 16 | 32 | 4 | >128 | 32 | 16 | 4 | 128 | 64 | 32 | >128 | 16 | 2 | 32 | 1 | 0.5 |
| Δ*mexB* Δ*mexF* | 0.5 | 2 | 0.5 | <0.125 | 1 | 1 | 0.5 | 4 | 4 | 8 | 4 | 4 | 16 | 2 | 16 | 1 | 0.5 |
| *mexR* ^PAO1^ | 4 | 4 | 4 | 0.5 | 64 | 16 | 8 | 4 | 64 | 8 | 8 | >128 | 16 | 2 | 32 | 1 | 0.5 |
| *mexT* ^PAO1^ | 64 | 16 | 32 | 4 | >128 | 16 | 4 | 2 | 128 | 64 | 32 | >128 | 16 | 2 | 32 | 1 | 0.5 |
| *mexR* ^PAO1^ *mexT* ^PAO1^ | 4 | 4 | 4 | 0.25 | 64 | 2 | 0.5 | 2 | 32 | 8 | 8 | 16 | 16 | 2 | 32 | 1 | 0.5 |
| *gyrA* ^PAO1^ | 64 | 16 | 32 | 4 | >128 | 2 | 4 | 4 | 128 | 64 | 32 | >128 | 16 | 2 | 32 | 1 | 0.5 |
| *gyrA* ^PAO1^ *mexR* ^PAO1^ | 4 | 4 | 4 | 0.5 | 64 | 2 | 0.5 | 4 | 64 | 8 | 8 | >128 | 16 | 2 | 32 | 1 | 0.5 |
| *gyrA* ^PAO1^ *mexT* ^PAO1^ | 32 | 16 | 32 | 4 | >128 | 2 | 0.5 | 2 | 128 | 64 | 32 | >128 | 16 | 2 | 32 | 1 | 0.5 |
| *gyrA* ^PAO1^ *mexR* ^PAO1^ *mexT* ^PAO1^ | 4 | 4 | 4 | 0.25 | 64 | 0.25 | <0.125 | 2 | 32 | 8 | 8 | 16 | 16 | 2 | 32 | 1 | 0.5 |
| *parS* ^PAO1^ | 64 | 16 | 32 | 4 | >128 | 32 | 16 | 4 | 128 | 64 | 32 | >128 | 16 | 2 | 32 | 1 | 0.5 |
| MIC breakpoint (intermediate) | 16 | 16 | 4 | 4 | 32 | 2 | 1 | 4 | - | - | - | - | - | 8 | - | 4 | 4 |
| **ATM**: Aztreonam; **CAZ**: Ceftazidime; **TZP**: piperacillin-tazobactam; **MEM:** Meropenem; **CAR**: Carbenicillin; **LVX**: Levofloxacin; **CIP**: Ciprofloxacin; **IPM**: Imipenem; **TMP**: Trimethoprim; **CRO**: Ceftriaxone; **TET**: Tetracycline; **CHL**: Chloramphenicol; **STR**: Streptomycin; **GEN**: Gentamicin; **FOF**: Fosfomycin; **PMB**: Polymyxin B; **CST**: Colistin | | | | | | | | | | | | | | | | | |
