## Supplementary material for "Native CRISPR-Cas mediated in situ genome editing reveals the exquisite interplay of resistance mutations in clinical multidrug resistant *Pseudomonas aeruginosa*": S3 Table

| Table S3: Contribution of basal-level, over-produced and overall MexAB-OprM pump to the resistance to different antipseudomonal antibiotics in PA154197  Different antibiotics | | | | | | | |
| --- | --- | --- | --- | --- | --- | --- | --- |
|  | ATM  (Monobactam) | CAZ  (Cephalosporin) | TZP  Penicillin-(β-lactamase) | MEM  (Carbapenem) | CAR  (Penicillin) | LVX  (Fluoroquinolone) | CIP  (Fluoroquinolone) |
| Relative contribution of basal-level MexAB-OprM  (MIC*_mexR_*^PAO1^/MIC_Δ_*_mexB_*) | 4 | 2 | 8 | >4 | 64 | 1 | 1 |
| Relative contribution of over-produced MexAB-OprM  (MIC_WT_/MIC*_mexR_*^PAO1^) | 16 | 4 | 8 | 8 | >2 | 2 | 2 |
| Overall contribution of efflux by MexAB-OprM  (MIC_WT_/MIC_Δ_*_mexB_*) | 64 | 8 | 64 | >32 | >128 | 2 | 2 |

Resistance contribution
