## Supplementary material for "Native CRISPR-Cas mediated in situ genome editing reveals the exquisite interplay of resistance mutations in clinical multidrug resistant *Pseudomonas aeruginosa*": S4 Table

| **Table S4.** Bacterial strains, plasmids used in this study | | | | |
| --- | --- | --- | --- | --- |
| **Strain or Plasmid No.** | **Description** | | | **Source** |
| Strains: |  |  | | |
| PA154197&PA150567 | *Pseudomonas aeruginosa* clinical isolate from Queen Mary Hospital (HK) | | From Prof. Patrick CY. WOO | |
| Ocean-100 | *Pseudomonas aeruginosa* strain isolated from the surface layer of the North Pacific Ocean | | | [1] |
| AY5236 | PA154197 Δ*mexB* | | | This study |
| AY5265 | PA154197 Δ*mexH* | | | This study |
| AY5266 | PA154197 Δ*mexF* | | | This study |
| AY5270 | PA154197Δ*mexB*&Δ*mexF* | | | This study |
| AY5912 | PA154197 *mexR*-T226G (*mexR* ^PAO1^) | | | This study |
| AY5929 | PA154197 *parS*-G1193A (*parS* ^PAO1^) | | | This study |
| AY5932 | PA154197 *mexT* ^PAO1^ | | | This study |
| AY5964 | PA154197 *gyrA*^PAO1^ | | | This study |
| AY6011 | PA154197 *mexT* ^PAO1^ & *mexR* ^PAO1^ | | | This study |
| AY6012 | PA154197 *gyrA*^PAO1^ & *mexR* ^PAO1^ | | | This study |
| AY6015 | PA154197 *gyrA*^PAO1^ & *mexT* ^PAO1^ | | | This study |
| AY6016 | PA154197 *gyrA*^PAO1^ & *mexT* ^PAO1^ & *mexR* ^PAO1^ | | | This study |
| Plasmids: |  | | |  |
| pMS402 | Reporter plasmid containing the promoter-less *lux* operon & *ori* of pRO1614 | | | [2] |
| pAY5211 | pMS402-P*tat* | | | This study |
| pAY5233 | pMS402-P*tat*-*mexB*(T) | | | This study |
| pAY5235 | pMS402-*mexB*(D-del)-P*tat*-*mexB*(T) | | | This study |
| pAY5244 | pMS402-P*tat*-*mexF*(T) | | | This study |
| pAY5245 | pMS402-*mexF*(D-del)-P*tat*-*mexF*(T) | | | This study |
| pAY5262 | pMS402-P*tat*-*mexH*(T) | | | This study |
| pAY5263 | pMS402-*mexH*(D-del)-P*tat*-*mexH*(T) | | | This study |
| pAY5901 | pMS402-P*tat*-*mexR*(T) | | | This study |
| pAY5902 | pMS402-P*tat*-*parS*(T) | | | This study |
| pAY5904 | pMS402-*mexR*(D-T226G)-P*tat*-*mexR*(T) | | | This study |
| pAY5905 | pMS402-*parS(*D-G1193A)-P*tat*-*parS*(T) | | | This study |
| pAY5919 | pMS402-P*tat*-*mexT*(T) | | | This study |
| pAY5920 | pMS402-*mexT*(D-1^st^ insertion)-P*tat*-*mexT*(T) | | | This study |
| pAY5923 | pMS402-P*tat*-(T-1^st^ insertion) | | | This study |
| pAY5925 | pMS402-*mexT*(D-PAO1)-P*tat*-(T-1^st^ insertion) | | | This study |
| pAY5949 | pMS402-P*tat*-*gyrA*(T) | | | This study |
| pAY5952 | pMS402-*gyrA*(D-PAO1)-P*tat*-(T-1^st^ insertion) | | | This study |
| **D**: donor; **T**: target |  | | |  |
